# SKiM-GPT: Combining Biomedical Literature-Based Discovery with Large Language Model Hypothesis Evaluation

**DOI:** 10.1101/2025.07.28.664797

**Authors:** Jack Freeman, Robert J. Millikin, Leo Xu, Ishaan Sharma, Bethany Moore, Cannon Lock, Kevin Shine George, Aviral Bal, Chitrasen Mohanty, Ron Stewart

**Affiliations:** Morgridge Institute for Research, Madison, WI 53715 USA

**Keywords:** literature-based discovery (LBD), large language models (LLM), retrieval- augmented generation (RAG), biomedical text mining, hypothesis evaluation, co-occurrence, fine-tuning

## Abstract

**Background:** Generating and testing hypotheses is a critical aspect of biomedical science. Typically, researchers generate hypotheses by carefully analyzing available information and making logical connections, which are then tested. The accelerating growth of biomedical literature makes it increasingly difficult to keep pace with connections between biological entities emerging across biomedical research. Recently developed automated means of generating hypotheses can generate many more hypotheses than can be easily tested. One such approach involves literature based discovery (LBD) systems such as Serial KinderMiner (SKiM), which surfaces putative *A B C* links derived from term co occurrence. However, LBD systems leave three critical gaps: (i) they find statistical associations, not biological relationships; (ii) they can produce false positive leads; and (iii) they do not assess agreement with a hypothesis in question. As a result, LBD search results often require costly manual curation to be of practical utility to the researcher. Large language models (LLMs) have the potential to automate much of this curation step, but standalone LLMs are hampered by hallucinations, lack of transparency in information sources, and the inability to reference data not included in the training corpus.

**Results:** We introduce **SKiM-GPT**, a retrieval-augmented generation (RAG) system that combines SKiM’s co-occurrence search and retrieval with frontier LLMs to evaluate user- defined hypotheses. For every chosen *A*-*B*-*C* SKiM hit, SKiM-GPT retrieves appropriate PubMed abstract texts, filters out irrelevant abstracts with a fine-tuned relevance model, and prompts an LLM to evaluate the user’s hypothesis, given the relevant abstracts. Importantly, the SKiM-GPT system is transparent and human-verifiable: it displays the retrieved abstracts, the hypothesis score, and a justification for the score grounded in the texts and written in natural language.

On a benchmark consisting of 14 disease-gene-drug hypotheses, SKiM-GPT achieves strong ordinal agreement with four expert biologists (Cohen’s κ = 0.84), demonstrating its ability to replicate expert judgment.

**Conclusions:** SKiM-GPT is open-source (https://github.com/stewart-lab/skimgpt) and available through a web interface (https://skim.morgridge.org), enabling both wet-lab and computational researchers to systematically and efficiently evaluate biomedical hypotheses at scale.

## Background

Literature-based discovery (LBD) is a field in which existing documented knowledge is searched, typically in an automated or semi-automated way, to aid in formulating hypotheses and finding novel discoveries by positing links between previously unconnected ideas (Smalheiser 2017). As the size of PubMed (currently over 37 million abstracts (Sayers et al. 2025)) and similar text databases grow, manual knowledge cataloging and connection-making becomes more difficult and LBD techniques become more important, as it is challenging for researchers to keep up to date with all relevant literature domains. Typically, an LBD “discovery” is in the form of an A-B-C relationship; A and B are two entities (genes, drugs, pathways, *etc.*) with an association, as are B and C. These two associations may imply a (possibly indirect) relationship between A and C, which may not yet be widely known. A classic example is fish oil as a treatment for Raynaud’s disease (Swanson 1987), discovered through their mutual link to blood viscosity.

A major advantage of automated LBD methods is their ability to rapidly perform hundreds or thousands of searches with different terms to identify latent associations at a scale that would be impractical manually (*e.g.*, with Google Scholar). However, while existing LBD methods can efficiently identify statistically significant associations between biomedical terms, they generally lack a structured framework for evaluating whether these discovered associations meaningfully support or contradict a specific scientific hypothesis. Tools such as Arrowsmith (Torvik and Smalheiser 2007), Lion-LBD (Pyysalo et al. 2019), BITOLA (Hristovski et al. 2006), and our tool Serial KinderMiner (SKiM) (Millikin et al. 2023) primarily quantify the strength of association without assessing relevance to or support of a hypothesis. See (Bhasuran et al., 2025) for a recent review of LBD methodologies. Consequently, researchers still face a significant burden of manually interpreting potentially thousands of associations, many of which are irrelevant or spurious (Preiss and Stevenson 2017). Thus, methods capable of automated, hypothesis-focused evaluation of LBD outputs are urgently needed.

Machine learning (ML), especially transformer-based large language models (LLMs), provide opportunities for contextualizing LBD-identified relationships due to their increased ability to capture nuanced semantic contexts compared to earlier approaches (Zhang et al., 2024; Zhao et al., 2025). Our previous augmentation of SKiM with ML-generated knowledge graphs showed promise but remained limited by sparse relationship annotations (Millikin et al., 2023). However, LLMs, particularly the advancement of reasoning models (J. Wei et al. 2023), are becoming increasingly capable of accurately parsing biomedical science texts. General purpose LLMs (OpenAI et al. 2024; Anil et al. 2024; DeepSeek-AI et al. 2025) achieve state□of□the□art scores on biomedical reasoning and question□answering benchmarks (Rein et al. 2023; Q. Chen et al. 2025; Jin et al. 2019). Furthermore, emerging multi-agent frameworks capable of autonomously generating and evaluating hypotheses (Gottweis et al. 2025) underscore a growing capability to not only surface potential discoveries but systematically assess their potential. Other methods for evaluation of hypotheses include benchmarking against curated datasets (Tyagin and Safro 2024) and utilizing knowledge graphs and transformer-based models (Tyagin et al. 2021). Motivated by these advances, we wanted to integrate LLM-based hypothesis evaluation into the SKiM workflow to enhance the interpretability and precision of putative LBD discoveries, while reducing the costs of manual review required of researchers.

However, a known challenge with LLMs is their propensity to generate incorrect or fabricated information (“hallucinations”) (Maynez et al. 2020). Retrieval-augmented generation (RAG) is a promising strategy to mitigate this issue by supplementing an LLM’s parametric (trained) memory with externally retrieved documents, providing a factual grounding for generated responses (Gao et al. 2024; Lewis et al. 2021). By combining both parametric (trained) and non-parametric (retrieved) sources, RAG reduces the frequency of hallucinations and improves interpretability, especially for information-sensitive tasks such as question answering and multi-hop reasoning (Pham and Vo 2024; Cao et al. 2023; Tonmoy et al. 2024).

Additionally, the recently developed scientific claim verification tool Valsci (Edelman & Skolnick, 2025) highlights the importance of RAG in minimizing the citation hallucination rate and reducing misclassifications of scientific claims as true or not. Our LBD tool, SKiM, serves two purposes. First, it is an effective way to surface new literature-based putative discoveries based on the co-occurrence model described above. Second, it provides a list of documents to be inputted for RAG analysis by the frontier LLM. These documents can supplement the model’s knowledge and provide context not available to the LLM during its training. Additionally, our RAG system, SKiM-GPT, displays these retrieved documents along with the LLM’s reasoning to the user, increasing transparency.

Still, the effectiveness of RAG depends on the quality and relevance of the retrieved documents. Retrieving documents that are irrelevant, ambiguous, or only weakly related to the hypothesis can introduce noise and misleading context, increasing the risk of inaccuracies and hallucinations (Hsieh et al. 2024; Modarressi et al. 2025). This risk is especially acute in biomedical literature, where terms frequently have multiple meanings and contexts can vary significantly (C.-H. Wei et al. 2019; Sohn et al. 2008). Therefore, a dedicated relevance filtering step is essential for RAG-based systems (J. Chen et al. 2024; Yu et al. 2024), ensuring that only information directly pertinent to the hypothesis at hand is included in the LLM’s context.

In summary, co-occurrence-based algorithms can discover novel associations without explicitly evaluating hypotheses, resulting in substantial manual effort to interpret relationships, manage high false-positive rates, and determine mechanistic context. LLMs have the potential to alleviate these problems but suffer from frequent hallucinations and opaque decision-making.

SKiM-GPT’s integrated approach leverages the complementary strengths of each tool, providing increased interpretability and precision compared to SKiM alone, while grounding the LLM’s reasoning in the retrieved documents. SKiM-GPT is open-source on GitHub (https://github.com/stewart-lab/skimgpt) and is available as a user-friendly web interface (https://skim.morgridge.org). Our contributions include: (1) providing a generalized method to improve the interpretability and false-positive rate of LBD systems; (2) providing an evaluation benchmark for that method; and (3) providing a concrete open-source implementation with a publicly accessible web interface.

### Implementation

The SKiM-GPT workflow is composed of three sequential steps: 1) identifying co-occurring terms in PubMed abstracts (“Serial KinderMiner”), 2) filtering abstracts for relevance to a user- defined hypothesis (“abstract filtering”), and 3) evaluating support for that hypothesis based on the filtered abstracts (“hypothesis evaluation”) (**Figure 1)**. Each of these steps is described below.

**Figure 1.**
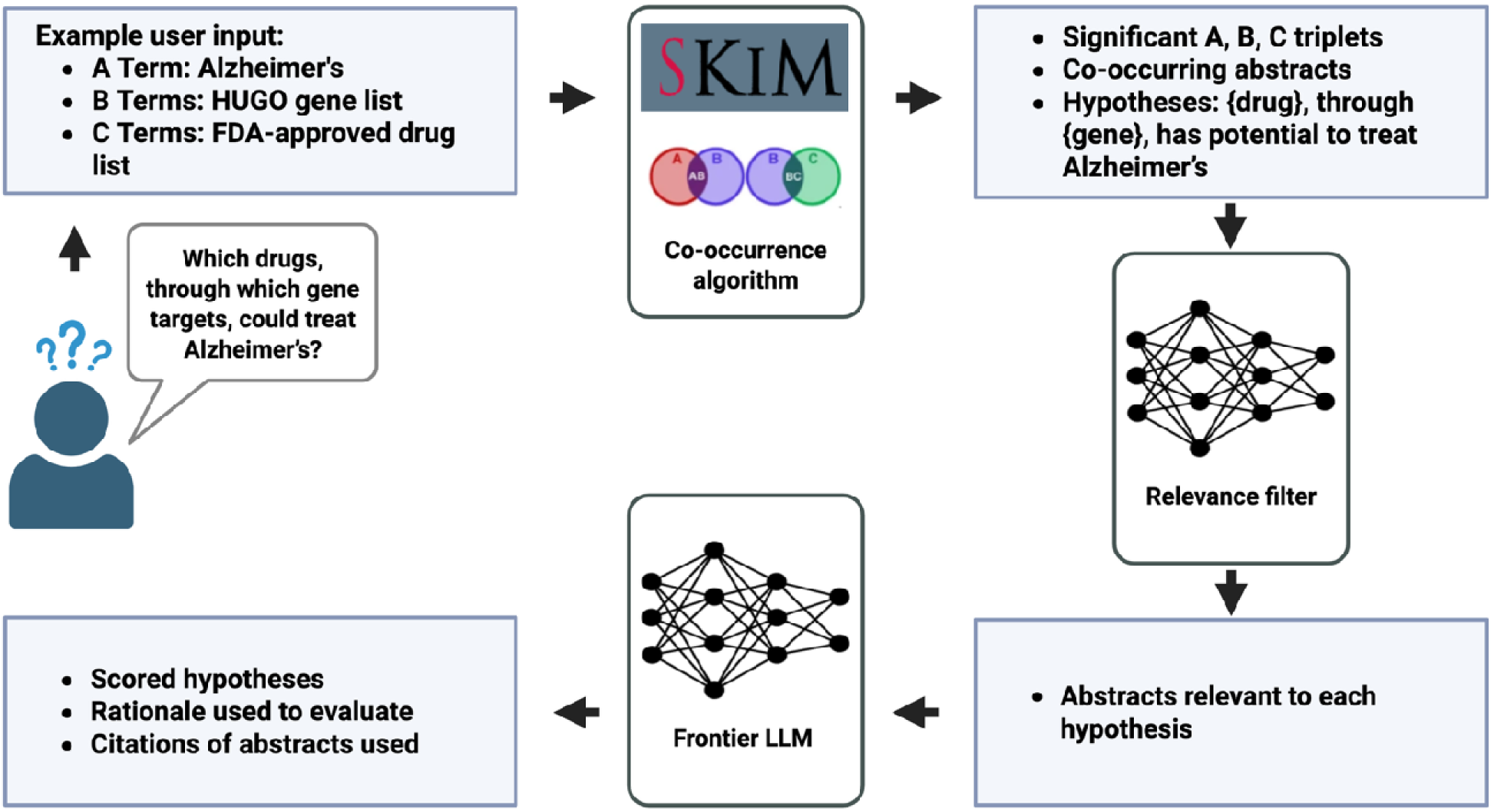
Overview of the SKiM-GPT workflow for hypothesis evaluation. Users input an A term and lists of B and C terms to discover hidden links between A and C terms. In this example, the user is interested in finding FDA-approved drugs that might be repurposed for Alzheimer’s disease, where one of the ∼19000 protein-coding human genes provide the link between drug and disease. SKiM is run to find potential A-B-C links. The A-B and B-C co-occurring abstracts found by SKiM are then run through the relevance filter to only include abstracts likely relevant to the hypotheses. Finally, these relevant abstracts and the hypotheses are evaluated by a frontier LLM, and the hypotheses are scored. The rationale and abstracts used are included in the output to provide full transparency.

#### Serial KinderMiner

The Serial KinderMiner (SKiM) algorithm is detailed in (Millikin et al. 2023). Briefly, SKiM identifies indirect relationships between A and C terms through B-term intermediates using co-occurrence modeling. Given user-defined lists of A, B, and C terms, SKiM searches PubMed abstracts for these terms using exact string matching and applies Fisher’s Exact Test (FET) to assess statistical significance of A-B and B-C term pairs using contingency tables of occurrence and co-occurrence counts (note that SKiM allows the use of “and” (&) and “or” (|) operators to search for inexact phrase matches and synonyms; this helps in cases when the user wants to search for pathways, biological processes, gene families and synonyms, etc. See the **Supporting Information** section “Exact String Matches and Evaluating Mechanism” for an example). A and C terms are considered putatively related if they share a significant B-term intermediary. SKiM ranks results based on a prediction score derived from *p*-values and term co-occurrence frequencies. In addition to statistical significance, abstracts containing co-occurring A-B, B-C, and A-C term pairs are retrieved as input for the next step.

#### Relevance filtering

After the SKiM search is complete, the user defines a hypothesis template into which the search terms can be inserted (*e.g.*, “Drug {c-term} treats disease {a-term} through its interaction with gene {b-term}.”). Abstract texts retrieved in the previous SKiM step are then filtered to determine their relevance to this hypothesis. Relevance is assessed using a fine-tuned language model based on the Phi-3 mini base architecture (Abdin et al. 2024), which outputs a single token indicating whether an abstract is relevant or irrelevant (“1” or “0”, respectively).

The primary goal of this step is to select abstracts that can be used to evaluate the hypothesis, regardless of whether they support or contradict it. This relevance filtering step is critical, as biomedical concepts frequently exhibit high ambiguity in free text (C.-H. Wei et al. 2019), and exact string-matching approaches like SKiM frequently retrieve abstracts containing ambiguous or irrelevant matches (Q. Chen et al. 2020). See **Supporting Information** section “Fine-Tuning Methodology” for further details on the implementation of relevance filtering.

#### Hypothesis evaluation

The hypothesis evaluation scoring guidelines, the user’s hypothesis, and the relevant abstracts are then processed by a frontier LLM. Inclusion of the relevant abstracts grounds the LLM’s analysis in the evidence contained in the abstracts rather than relying on its parametric memory. The LLM assigns a numerical score between -2 and 2, representing the degree of support for the hypothesis according to the scoring guidelines, and provides explanatory reasoning including references to the abstracts used to arrive at a score. For scoring guidelines and an example prompt see **Supporting Information.**

#### SKiM-GPT web API server implementation

SKiM-GPT was implemented as an open-source Python package (https://github.com/stewart-lab/skimgpt) and incorporated into the existing SKiM web API server (Millikin et al., 2023) as a new job type. Briefly, the SKiM web API server uses a job queue system in which jobs are submitted to the server via HTTP request and are put into the work queue. Workers retrieve the jobs and do the requisite processing (in this case, using the SKiM-GPT python package to perform the relevance filtering and hypothesis evaluation steps). The job’s status and results can be retrieved via HTTP request. The server is intended to be run as a multi-container Docker application; Python code and Docker build scripts can be found at https://github.com/stewart-lab/fast_km. A public website for running SKiM and SKiM-GPT queries that uses this API can be found at https://skim.morgridge.org.

#### Score alignment evaluation

Fourteen disease-gene-drug combinations were chosen from statistically significant SKiM hits (twelve by RM; two by RS), intended to represent a wide variety of scores (i.e., -2 to +2). Two of these examples were chosen because they were false-positive SKiM hits (pancreatic cancer-CCK-gant61, diabetes-AHR-stauvidine). The rationale for why these examples were chosen was not shared with the other co-authors until the scoring was completed. The primary motivation behind these choices was to provide a set of examples to assess how the system performed under relatively diverse scoring conditions.

## Results

### Score Alignment Evaluation

To assess how SKiM-GPT scores align with those of human evaluators, we gave four expert computational biologists (JF, RM, IS, RS) the task of evaluating a series of drug repurposing hypotheses and then compared those human scores to SKiM-GPT’s. Fourteen distinct disease-gene-drug combinations were generated for this task from statistically significant SKiM hits. Human and LLM evaluators scored the supporting abstracts on a scale from −2 (strongly refutes the hypothesis) to +2 (strongly supports the hypothesis). SKiM-GPT generated scores from 10 independent runs per hypothesis to get a score distribution for each hypothesis (see **Figure 2** for an example output).

**Figure 2.**
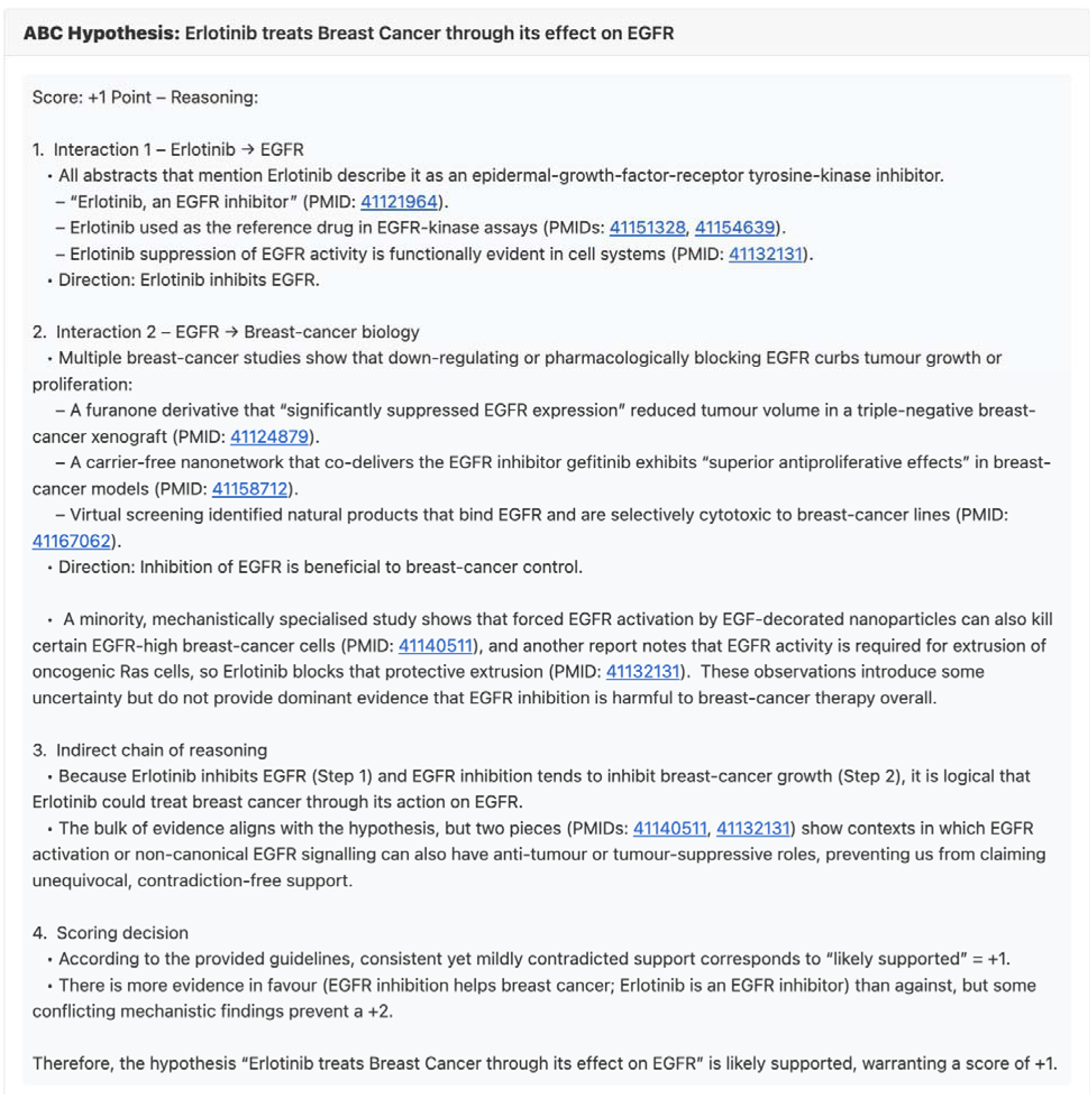
Screenshot of the web interface after evaluating the A-B-C hypothesis “Erlotinib treats Breast Cancer through its effect on EGFR”.

Without relevance filtering, SKiM-GPT showed substantial agreement with human evaluators (**Figure 3A**; root mean square error (RMSE) = 0.57, quadratic-weighted Cohen’s_κ_ (QWK) = 0.84 (Fleiss and Cohen 1973), intraclass correlation coefficient (ICC (2,1)) = 0.91 (Koo & Li, 2016). Directional concordance between SKiM-GPT’s median score and human median score occurred in 13 out of 14 hypotheses (93%), and the median SKiM-GPT score fell within the human score range in 12 of 14 cases (79%).

**Figure 3.**
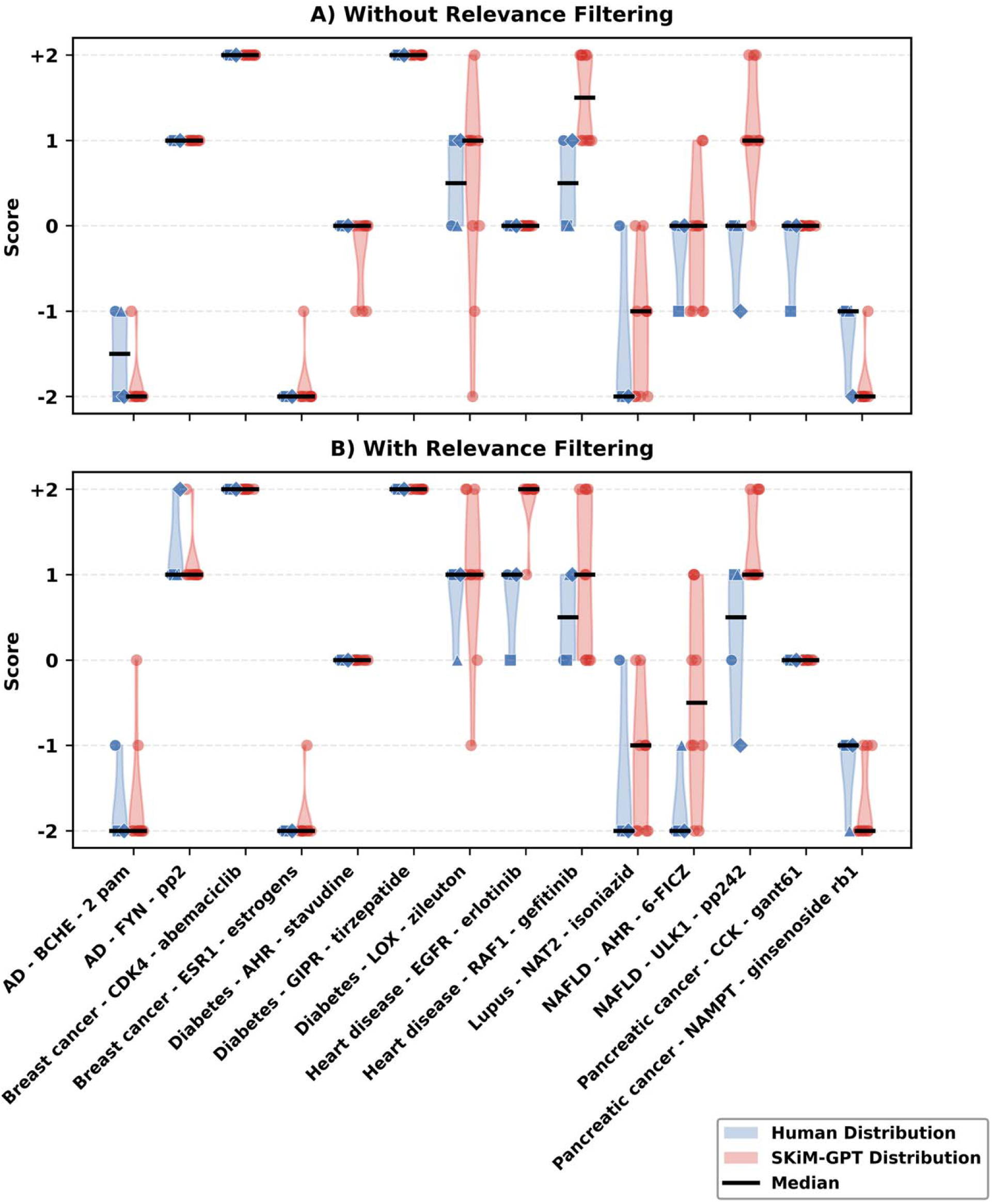
Comparison between human expert and SKiM-GPT hypothesis evaluation scores across 14 disease–gene–drug hypotheses. The hypotheses were scored in a range from −2 (strongly refuted) to +2 (strongly supported) based on the provided abstracts. **(A)** *Without relevance filtering.* For each hypothesis, red violins show the distribution of SKiM-GPT scores obtained from 10 independent runs; overlaid red dots are the individual run values. Blue violins show the distribution of the four expert scores, with overlaid blue symbols marking each evaluator’s score. Horizontal lines inside violins mark medians. Median SKiM-GPT scores largely agreed with human evaluators (directional concordance = 93%; within-range agreement = 86%; RMSE = 0.57). In cases where the distributions are narrow, there is generally little subjective interpretation needed; the score is fairly obvious. In less obvious cases, the distributions are wider because the scores are more subjective or uncertain. **(B)** *With relevance filtering.* Same plotting conventions as in (A). With relevance filtering, SKiM-GPT median scores maintained strong agreement with human evaluators (directional concordance = 100%, within-range agreement = 86%; RMSE = 0.64) compared to no relevance filtering.

A relevance filtering step was added to ensure that human and SKiM-GPT evaluators based their decisions on abstracts directly pertinent to the hypotheses (see **Relevance Filtering Evaluation** below). This filtering step removes irrelevant or contextually ambiguous abstracts retrieved by SKiM (*e.g.*, “AHR” can refer to the “aryl hydrocarbon receptor” gene or the “adjusted hazard ratio” measure). With relevance filtering, agreement between SKiM-GPT and human evaluators remained high (**Figure 3B**; RMSE = 0.64, QWK= 0.84, ICC (2,1) = 0.91), with directional concordance of 100% (14/14) and median scores falling within the human evaluators’ range in 12 of 14 cases (86%).

### Relevance Filtering Evaluation

To evaluate relevance classification performance, we compared the pretrained base and fine-tuned models using a set of 177 abstract–hypothesis pairs (140 manually annotated from Figure 3 literature and 37 LLM-generated [“synthetic”] abstracts). Each pair was labeled as either 0 (“Not Relevant”) or 1 (“Relevant”) and split into 105 training and 72 test pairs. The pretrained model achieved 0.68 accuracy, 0.70 precision, 0.92 recall, and an F1 of 0.80, while the fine-tuned model improved performance to 0.86 accuracy, 0.85 precision, 0.96 recall, and an F1 of 0.90 (Figure 4A–B). The fine-tuned model misclassified only two relevant abstracts as irrelevant, reflecting a deliberate design choice to favor a low false-negative rate so that nearly all potentially useful evidence is retained for downstream hypothesis evaluation. Next, we measured the model’s performance on an out-of-distribution abstract set to assess the generalizability of the relevance filter to hypotheses not observed during training. In this evaluation, abstracts containing possible drug-drug interactions (DDIs) (**Figure 4C**) and functional gene interactions (FGIs) (**Figure 4D**) were generated with GPT-4 and labeled with a 1 or 0, as above. The model’s precision 0.90 and recall 1.0 on this out-of-training data indicates the ability to generalize across biomedical domains. For more information on generating synthetic text and descriptions of relevance categories, see **Supporting Information** and **Figures S1A-B.**

**Figure 4.**
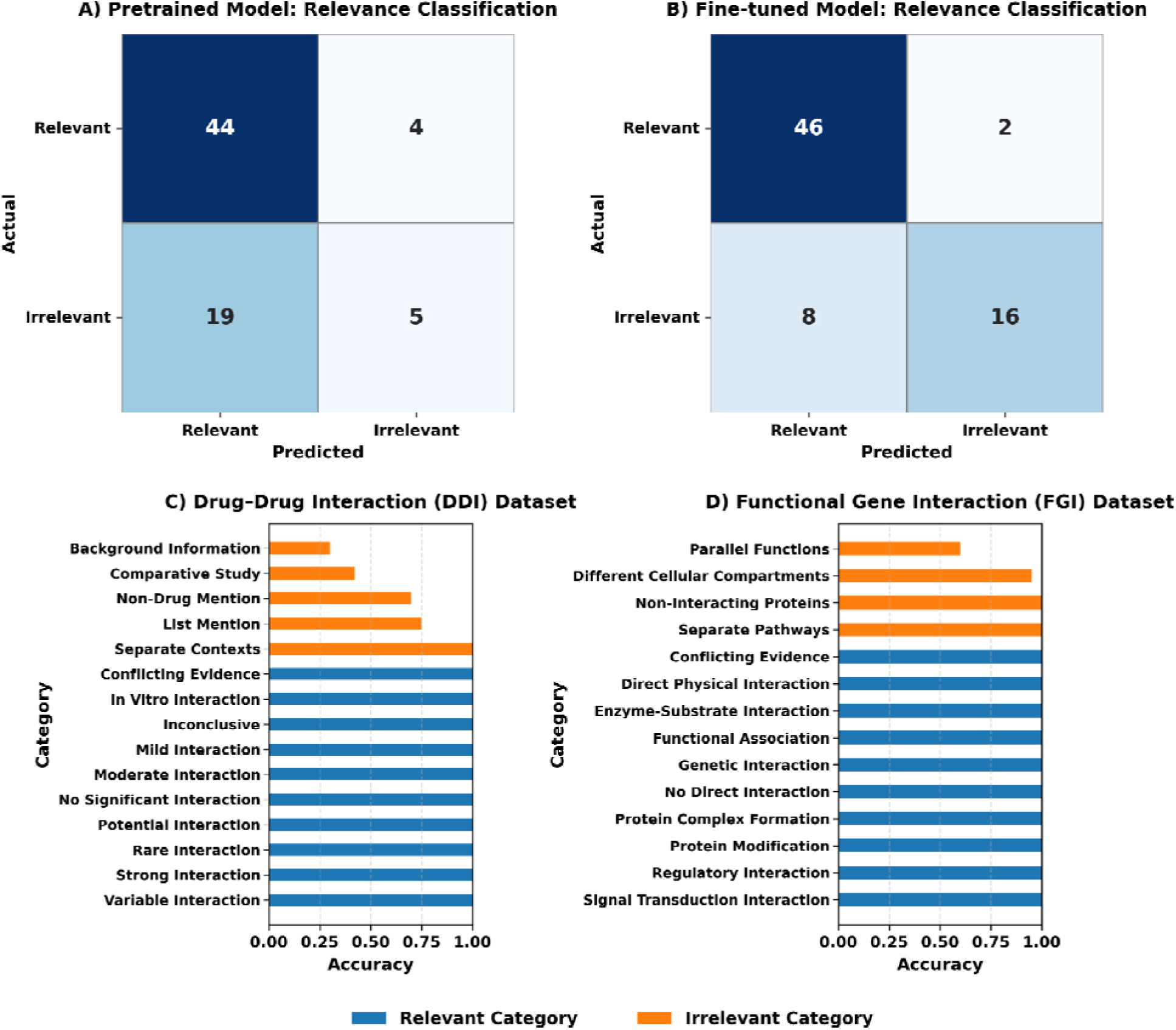
Evaluation of the relevance-filtering model. (**A**) *Pretrained base model (Phi-3-mini- 4k-instruct) performance confusion matrix.* Confusion matrix showing performance of the pretrained base model on the 72-abstract test set. The model correctly retained 44 of 48 relevant abstracts and correctly discarded 5 of 24 irrelevant abstracts (accuracy = 0.68; precision = 0.70; recall = 0.92; F1 = 0.80). **(B)** *Fine-tuned model performance confusion matrix.* Evaluation of the relevance filtering model on the 72-abstract test set shows that 46 of 48 relevant abstracts were correctly retained, while 16 of 24 irrelevant abstracts were correctly discarded (accuracy = 0.86; precision = 0.85; recall = 0.96; F1 = 0.90). Two relevant abstracts were incorrectly discarded, and eight irrelevant abstracts were incorrectly retained, reflecting the design choice to favor low false-negative rates so that almost all useful evidence reaches the hypothesis evaluation stage. **(C, D)** *Generalization to unseen interaction types.* Bar charts report per-category accuracy on 100 GPT-4-generated abstracts describing drug–drug interactions **(C)** and functional gene interactions (**D**) that were absent from training. Blue bars indicate contexts expected to be relevant to the hypothesis; orange bars indicate background or non-interaction scenarios that should be rejected as irrelevant.

### Cost Estimation

One way to understand the discrepancy between the vast amount of published biomedical literature and the human bandwidth available to read and interpret it is through an analysis of the cost difference between human and SKiM-GPT evaluation. Human evaluators required an average of 8.42 ± 1.20 hours per person to read 159 abstracts, incurring an estimated cost of $336.80 per person (assuming $40/hour) (**Table 1**). In contrast, the SKiM-GPT system evaluated the same set of abstracts in 0.23 ± 0.03 hours, with a cost of $8.62 (the OpenAI API call cost). This represents a 97% reduction in evaluation time and a 97% decrease in monetary cost when using SKiM-GPT compared to manual human analysis.

**Table 1.** Cost difference between human and SKiM-GPT evaluation.

| Number of Abstracts | Human Evaluation Time (hours) | SKiM-GPT Evaluation Time (hours) | Human Evaluation Monetary Cost (USD) | SKiM-GPT Evaluation Monetary Cost (USD) |
| --- | --- | --- | --- | --- |
| 159 | $8.42 \pm 1.20$ | $0.23 \pm 0.03$ | \$336.80 | \$8.62 |

**Table 1**. Cost estimation for human experts and SKiM-GPT to evaluate 159 biomedical abstracts. SKiM-GPT took 0.23 ± 0.03 hours to review the given text corpus at a cost of $8.62 in API calls, whereas human experts required 8.42 ± 1.20 hours (est. $336.80, assuming $40/hour). SKiM-GPT was 97% faster and 97% less expensive compared to human experts.

### Retrieved Text vs Memory

We next assessed the potential problem of parametric-knowledge leakage in SKiM-GPT’s evaluations through a series of controlled experiments. “Parametric-knowledge leakage” refers to the LLM’s undesired reliance on internal parametric memory to evaluate a hypothesis instead of external information (*i.e.*, the abstracts). The desired behavior for SKiM-GPT’s hypothesis evaluation is to rely only on the provided, contextually relevant literature, rather than pre-existing model biases. To assess leakage, we had SKiM-GPT evaluate three different hypotheses that assess the efficacy of FDA-approved breast cancer drugs (**Figure 5A**) in improving breast cancer patient outcomes. Synthetic (*i.e.*, fake) abstracts were generated to imply neutral, negative, and positive relationships between each drug and breast cancer patient outcomes and supplied to SKiM-GPT. When the positive hypothesis was evaluated without any retrieved abstracts, SKiM-GPT produced strongly positive scores, confirming expected leakage from the training data (**Figure 5B**). SKiM-GPT strongly supported neutral hypotheses when provided with neutral abstracts (**Figure 5C**), negative hypotheses when supplied with negative abstracts (**Figure 5D**), and positive hypotheses when provided with positive abstracts (**Figure 5B**), indicating that SKiM-GPT grounds its analysis in the provided texts even in cases where its parametric memory conflicts with those texts **(Figure 5C and 5D)**. When using the five most cited real abstracts, the positive scores indicated correct agreement with the positive hypothesis (**Figure 5B**). Additionally, we evaluated parametric-knowledge leakage using real abstracts; two examples are provided in the **Supporting Information** section “Time-Slicing Parametric- Knowledge Leakage Evaluation”.

**Figure 5.**
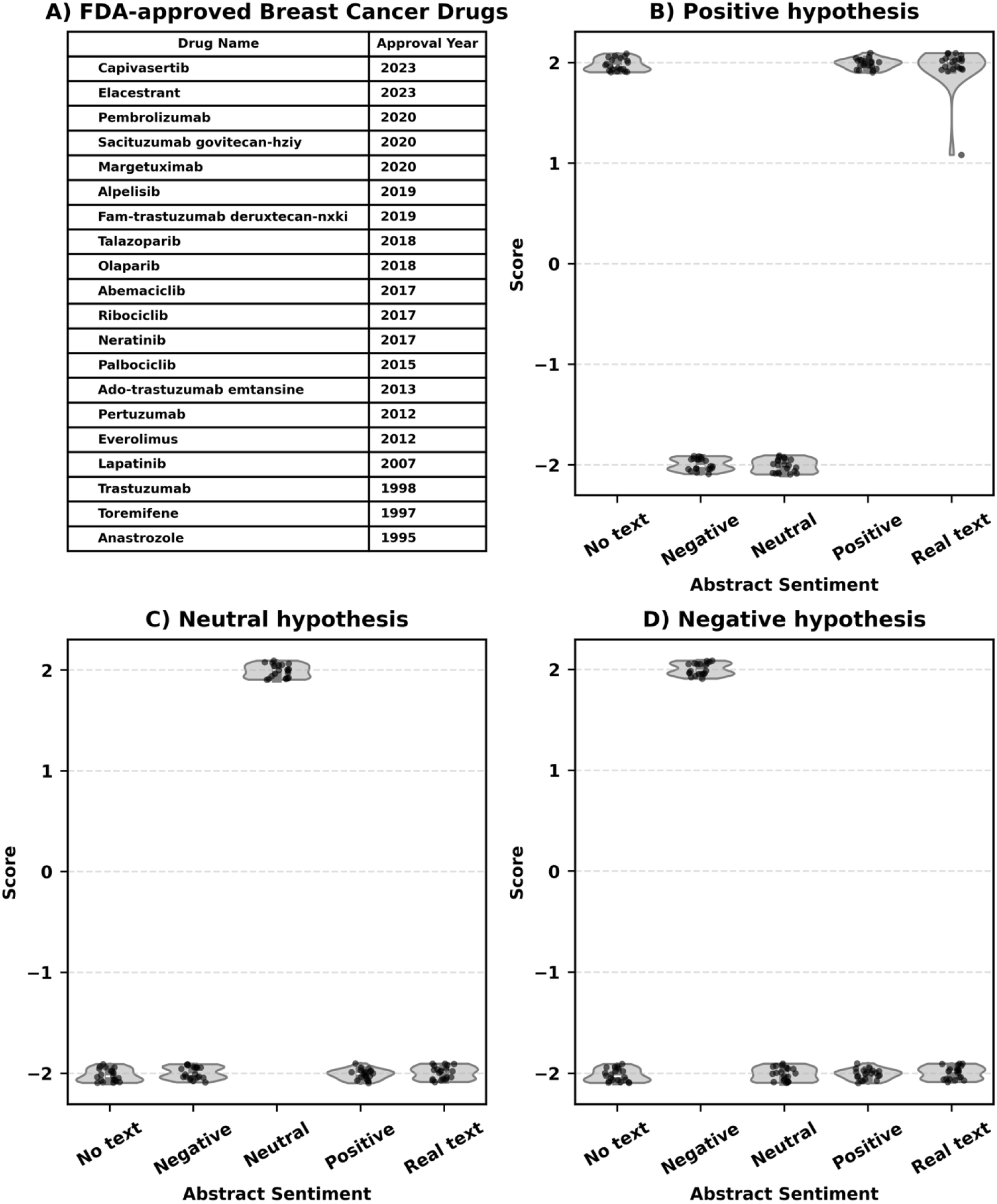
Parametric-knowledge leakage test of SKiM-GPT on 20 FDA-approved breast cancer drugs. **(A)** Reference table listing each drug and its FDA approval year. **(B-D)** Each point represents the hypothesis support score for each drug. Violin plots with y-axis jittered points depict score distributions across conditions on the x-axis, where “Negative,” “Neutral,” and “Positive” denote synthetic abstract sentiment and “Real text” denotes retrieved PubMed abstracts while “No text” denotes no added abstracts. **(B)** *Positive hypothesis condition.* Without any provided abstracts (“No text”), when asked whether each drug “improves breast-cancer patient outcomes,” SKiM-GPT assigns positive scores (median = +2), indicating that parametric (trained) memory is biased towards positive scores in these cases as expected (“parametric- knowledge leakage”). **(C)** *Neutral hypothesis condition.* Feeding synthetic abstracts that state no relationship exists between the drug and breast cancer patient outcomes increases scores for neutral hypotheses (median = +2) for all drugs, indicating the model grounds its judgment in the supplied context rather than internal bias. **(D)** *Negative hypothesis condition.* Providing synthetic abstracts that refute drug efficacy results in increased scores for negative hypotheses (median = +2). Together, panels B–D indicate that SKiM-GPT’s scores mirror the sentiment of the abstracts rather than the LLM’s training data.

### LBD False-Positive Removal Evaluation

Co-occurrence-based LBD search systems like SKiM measure how frequently search terms occur together in a corpus of text. These frequency measures (i.e. *p*-values) do not describe how or why these terms co-occur. The addition of relevance filtering and frontier LLM hypothesis evaluation allows ranking of true-positives in the co-occurrence-based LBD hit list, and removal of false-positives. Above, we have discussed examples where relevance filtering removes false-positive hits owing to semantic ambiguity (*e.g.*, the “AHR” search term can refer to “aryl hydrocarbon receptor” or “adjusted hazard ratio”). In these cases, false-positive abstracts are removed via relevance filtering. In other cases, SKiM may primarily retrieve abstracts in which the terms correctly refer to the user’s intended meaning, but the retrieved abstract texts disagree with the user’s hypothesis and is thus a different type of false-positive LBD hit.

To test whether SKiM-GPT can remove false-positives and rank true-positives, we queried SKiM using “breast cancer” as the A-term, a comprehensive list of human genes as B- terms, and a comprehensive list of FDA-approved drugs as C-terms. The hypothesis used for evaluation was “*{C-term} treats {A-term} through its effect on {B-term}.*” The top 20 highest-ranked A-B-C hits by *p*-value (see **Supporting Information** for methodological details) are reported in **Table 2**. Manual evaluation of these top 20 hits indicated that SKiM-GPT correctly classified 19 of them, confirming 16 true drug-gene treatment links (positive scores) and correctly discarding three false positives (zero or negative scores). In the one case where SKiM- GPT was unable to correctly classify, the gene name SLN returned no relevant abstracts, as the abbreviation SLN can also stand for “sentinel lymph node” which is often a term associated with cancers.

**Table 2.** SKiM-GPT’s ability to filter a list of SKiM co-occurrence hits.

| A-B-C Relationship | Score | Label | Comments |
| --- | --- | --- | --- |
| Breast Cancer - BCL2 - VENETOCLAX | +1 | TP |  |
| Breast Cancer - BRCA1 - OLAPARIB | +2 | TP |  |
| Breast Cancer - BRCA2 - OLAPARIB | +2 | TP |  |
| Breast Cancer - CDK4 - ABEMACICLIB | +2 | TP |  |
| Breast Cancer - CDK4 - FULVESTRANT | -1 | TN | FULVESTRANT doesn't treat breast cancer via CDK4 directly. |
| Breast Cancer - CDK4 - PALBOCICLIB | +2 | TP |  |
| Breast Cancer - EGFR - AFATINIB | +1 | TP | Only mildly effective in some breast cancer cases |
| Breast Cancer - EGFR - CETUXIMAB | +1 | TP |  |
| Breast Cancer - EGFR - CRIZOTINIB | -2 | TN | CRIZOTINIB doesn't act directly through EGFR and is not known to have indirect effects on the EGFR pathway. It may be able to combine with EGFR inhibitors to treat breast cancer |
| Breast Cancer - EGFR - DACOMITINIB | +1 | TP | Not a primary treatment. Sometimes used when other treatments fail |
| Breast Cancer - EGFR - ERLOTINIB | +1 | TP | Only effective for some breast cancer patients |
| Breast Cancer - EGFR - GEFITINIB | +1 | TP | Only effective for some breast cancer patients |
| Breast Cancer - EGFR - PANITUMUMAB | +1 | TP | PANITUMUMAB can treat aggressive forms of inflammatory breast cancer via EGFR inhibition. |
| Breast Cancer - EGFR - PEMETREXED | -1 | TN | Plausible mechanism but mostly used for NSCLC |
| Breast Cancer - ESR1 - FULVESTRANT | +1 | TP |  |
| Breast Cancer - MTOR - EVEROLIMUS | +2 | TP |  |
| Breast Cancer - MTOR - TEMSIROLIMUS | +2 | TP |  |
| Breast Cancer - PIK3CA - ALPELISIB | +1 | TP |  |
| Breast Cancer - RAD51 - OLAPARIB | +1 | TP | While OLAPARIB doesn't act directly via RAD51, RAD51 deficiency allows OLAPARIB's effectiveness |
| Breast Cancer - SLN - ISOSULFAN BLUE | N/A | False N/A | No relevant abstracts were found; should get a negative score because ISOSULFAN BLUE doesn't treat breast cancer via SLN gene. |

**Table 2**. Assessment of SKiM-GPT’s ability to remove false-positive hits in an LBD search. “Breast Cancer” was used as the A-term, a list of human genes as B-terms, and FDA-approved drugs as C-terms. The hypothesis “*{C-term} treats {A-term} through its effect on {B-term}*” was used to evaluate the top 20 hits ranked by *p*-value. For each SKiM A-B-C hit (column 1), SKiM- GPT scores the provided hypothesis from –2 (strongly refutes) to +2 (strongly supports) after relevance filtering (column 2). Scores were classified as a true-positives (TP) or true-negatives (TN) based on current clinical or mechanistic evidence (column 3). The comments (column 4) contain context-specific notes. SKiM-GPT correctly classified 19 of 20 hits, confirming 16 valid drug–gene treatment links and correctly discarding three false positives.

## Discussion

Recent advances in natural language processing have enabled increasingly context-aware automated parsing of the biomedical literature at scale, rather than relying on simple text search and manual validation. SKiM-GPT pairs SKiM’s co-occurrence search and retrieval with a relevance filter and an LLM-based hypothesis evaluator. This three-stage workflow discards many of the spurious associations as documented in earlier LBD studies (Gopalakrishnan et al. 2019; Cheerkoot-Jalim and Khedo 2021), anchors each evaluation in the provided literature, and produces quantitative and qualitative support that aligns with expert judgement. While earlier tools such as KinderMiner and SKiM excel at finding associations between entities by surfacing statistically enriched term pairs (Kuusisto et al. 2017; Millikin et al. 2023), SKiM GPT defines relationships and provides the precision that co occurrence models lack, without exposing it to the hallucinations and other issues such as lack of transparency, opaqueness of reasoning, and parametric-knowledge leakage that plague stand alone LLMs.

The platform is freely accessible as both a command line utility and a web application (https://skim.morgridge.org/), with the web interface meant to be usable by anyone at any skill level. Some example goals of this system are to 1) discover new, potentially meaningful connections between biomedical concepts and assess their plausibility for downstream experimental validation, 2) evaluate a new hypothesis that the user has, 3) re-examine or benchmark existing hypotheses using time-sliced or current literature evidence. Some examples of potential SKiM-GPT uses include but are not limited to prioritizing drug repurposing candidates, interrogating functional gene or drug–drug interactions, assembling gene regulatory networks, identifying pathways that link genes, and/or refining cell type annotations in single cell atlases without manually scouring the literature. While many associations are likely established in literature, SKiM-GPT may also find novel A-B-C links that can lead to new hypotheses and discoveries. For more information on identifying novel A-B-C links see **Supporting information** section “How to Identify Novel Hypotheses Returned by SKiM-GPT*”*.

At present SKiM GPT analyzes only PubMed titles and abstracts, leaving valuable insights in full-text articles, figures, and tables inaccessible. It also lacks support for non-textual modalities such as microscopy images, clustering plots, and single-cell embeddings. Moreover, large-scale screens can be expensive due to reliance on commercial LLM APIs. To address these limitations, we plan to implement an open-source hypothesis evaluator model, eliminating dependence on proprietary models and their API costs. We also plan to extend retrieval beyond abstracts to full-text articles and other structured data, and to reduce future inference costs through model quantization and routing techniques. Additionally, negative or inconclusive results are less likely to be published, and these unpublished results would be invisible to SKiM- GPT; this may skew SKiM-GPT’s scores, especially in the case where few abstracts are available to score a hypothesis. Regularization or a Bayesian prior may help address this issue.

We would also like to modify the relevance filter to rank the retrieved texts by a continuous relevance score and taking the top N with some minimum relevance score, rather than using a binary classifier for relevant/irrelevant and taking N. This may help make some hypotheses easier to score; some examples of difficult-to-score hypotheses in **Figure 3** could be a good starting point for fine-tuning this new relevance filtering model.

SKiM GPT couples high recall retrieval with context aware LLM reasoning to surface the most plausible mechanistic relationships hidden in an ever growing corpus of biomedical literature, making hypothesis evaluation 97% faster and 97% less expensive for both dry and wet lab researchers.

## Availability and Requirements

Project Name: SKiM-GPT.

Project home page: https://skim.morgridge.org/ (user-facing website); https://github.com/stewart-lab/skimgpt (application code).

Operating system(s): Platform independent.

## Programming language: Python

Other requirements: To run SKiM-GPT from the command line, Python 3.10 or higher is required along with other requirements outlined in requirements.txt. Additionally, a user must have an account with OSPool or other HTCondor systems with GPU access; to use the tool on the website, no requirements except a web browser.

## License: MIT

Any restrictions to use by non-academics: No restrictions.

### List of abbreviations

RMSE: Root mean square error
ICC: Intraclass correlation coefficient
LLM: Large language model
RAG: Retrieval-augmented generation
LBD: Literature-based discovery
KM: KinderMiner (two-term co-occurrence search)
SKiM: Serial KinderMiner (A-B-C three-term co-occurrence framework)
GPT: Generative pre-trained transformer
HTTP: Hypertext transfer protocol
API: Application programming interface
DDI: Drug-drug interaction
FGI: Functional gene interaction
QWK: Quadratic weighted kappa
FET: Fisher’s exact test

## Declarations

### Ethics approval and consent to participate

**Not applicable.** This work analyzed publicly available biomedical abstracts and did not involve human participants.

## Consent for publication

**Not applicable.** The manuscript does not contain any individual person’s data.

## Availability of data and materials

The SKiM-GPT package is open-source on GitHub (https://github.com/stewart-lab/skimgpt) All synthetic text data, term lists, and relevance filter misclassifications are available (https://github.com/stewart-lab/skimgpt/tree/paper) This data is also included in the additional file “SKiM_GPT_supplementary_Data.zip”.

## Authors’ Contributions

JF, LX, KG, RM, and RS developed SKiM-GPT. JF, RM, and RS designed the study and evaluations. JF, RM, IS, and RS evaluated the hypotheses. LX and JF trained and integrated the relevance filter. JF and IS created the synthetic datasets. BM, JF and RM provided statistical support. CL, AB, KG, RM and JF built the web interface. CM ran the dimensionality reduction experiment. JF, RM, IS and RS wrote the manuscript.

## Acknowledgements

We thank the Center for High Throughput Computing (CHTC) at UW–Madison for GPU resources and the Morgridge Institute for Research Computing staff for infrastructure support. We thank Alicia Williams for editorial assistance.

