## Supporting Information for "SKiM-GPT: Combining Biomedical Literature-Based Discovery with Large Language Model Hypothesis Evaluation"

### **Title**

###### Generating and Evaluating Synthetic Abstracts

For both fine-tuning and parametric leakage experiments, we constructed sets of synthetic biomedical abstracts through an open-source generation pipeline **(** <https://github.com/stewart-lab/skimgpt/blob/paper/paper/abstract_generator.py> **)**. We developed a Python-based generation framework that interfaces with either the OpenAI GPT-4 API or the Deepseek API via the “openai” Python package. For bias control, we intentionally alternated model families between *generation* and *evaluation*: during fine-tuning when the fine-tuned *Phi-3-mini-4k-instruct* relevance model served as the evaluator, abstracts were generated with GPT-4; in the parametric leakage experiments *o1* performed the evaluation, thus abstracts were generated with DeepSeek-R1. This cross-family pairing reduced the risk of inadvertent alignment between generation and evaluation models due to overlapping linguistic priors. Additionally, for synthetic abstracts, PMIDs are replaced with randomly generated placeholders (e.g., “PMID: [RANDOM NUMBER]”) for format consistency. These do not correspond to real PubMed records. For specifics in developing each set of abstracts for either fine-tuning the relevance model or testing parametric leakage, please see the corresponding section.

To assess whether our synthetic abstracts are representative of authentic ones, we embedded 100 real abstracts (five per drug) alongside 200 synthetic abstracts (five positive-sentiment and five negative-sentiment per drug) spanning 20 drugs. These are the same abstracts that were used in our parametric-knowledge leakage experiment **(Figure 5)**.

We then applied three dimensionality reduction techniques, PCA, t-SNE, and UMAP, to visualize how the embeddings cluster. The resulting projections show substantial overlap between real and synthetic abstracts (**Figure S1A**). When colored by drug, clustering is driven primarily by drug identity rather than by real or synthetic origin: for each drug, the real and synthetic abstracts co-locate, and within-drug distances are smaller than between-drug distances (**Figure S1B**). These results suggest that large language models encode synthetic and real abstracts in a similar manner.

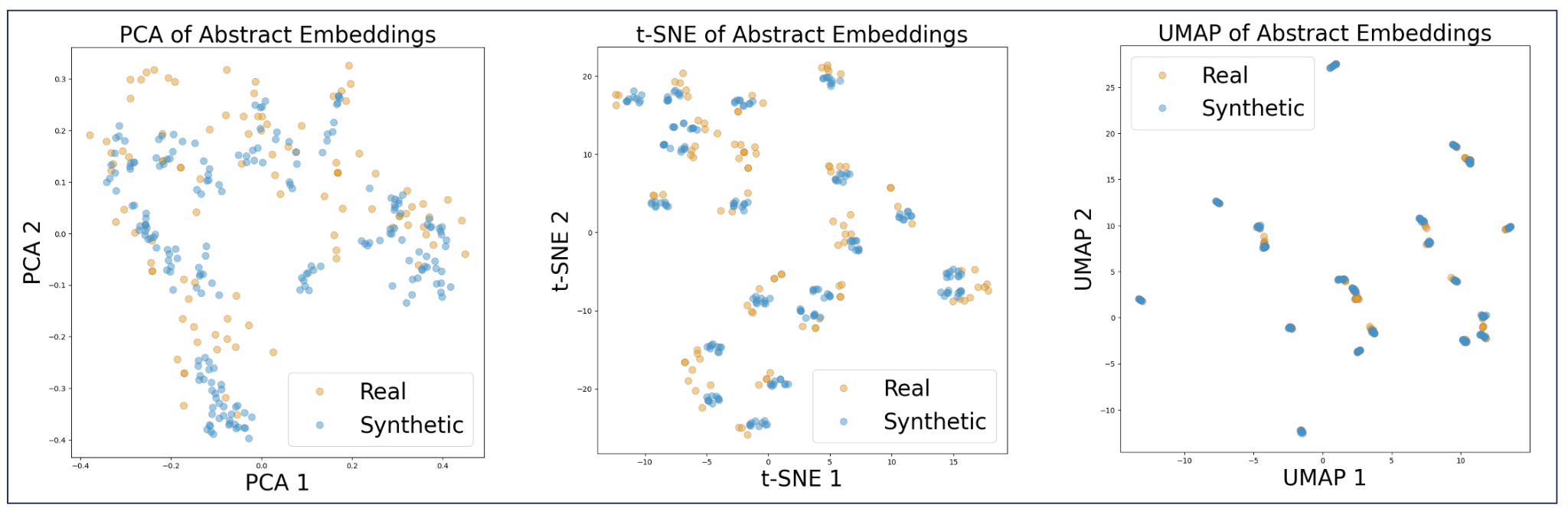

**Figure S1A:** Two-dimensional visualizations of the embeddings from real and synthetic abstracts using PCA, t-SNE, and UMAP.

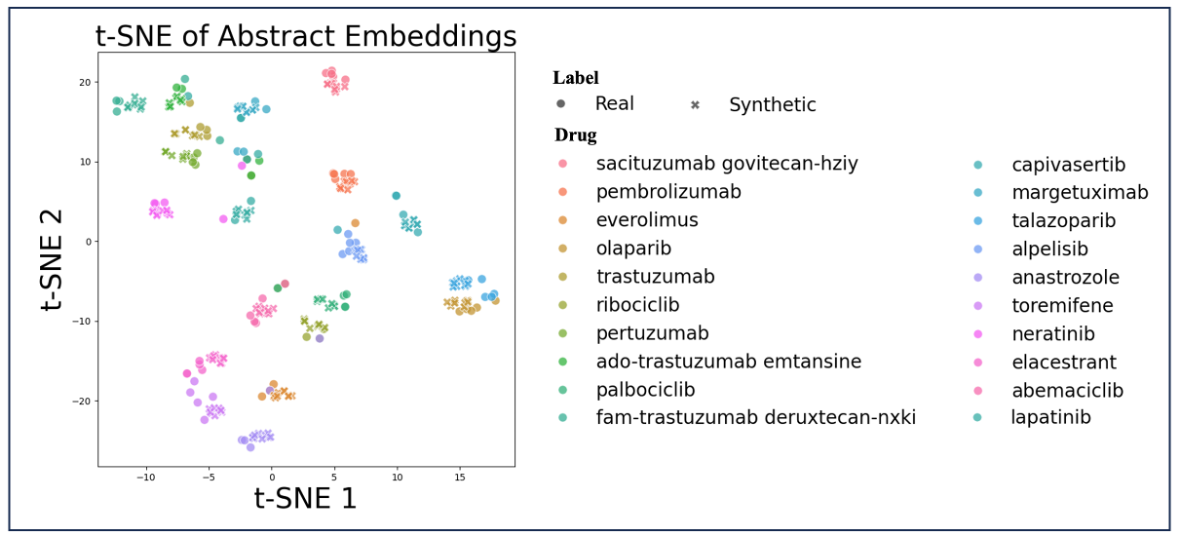

**Figure S1B**: Two-dimensional visualization of the embeddings using t-SNE and colored by drug. Real and synthetic abstracts that mention a particular drug cluster together indicating that the difference between synthetic and real abstracts that mention the same drug is less than the difference between synthetic abstracts that mention different drugs or real abstracts that mention different drugs.

###### Fine-Tuning Methodology

The relevance filter was implemented using the unsloth/Phi-3-mini-4k-instruct model, (Unsloth, 2025) fine-tuned with a sequence length of 2048 tokens. Rank-Stabilized Low-Rank Adaptation (rsLoRA) (Hu et al., 2021; Kalajdzievski, 2023) parameters were configured with rank (r) set to 16, LoRA alpha at 32, and no dropout. Additionally, during training, Noise-Enhanced Fine-Tuning (NEFTune) was applied (Jain et al., 2023). To enhance computational efficiency, flash_attention_2 (Dao, 2023) was utilized as the attention mechanism. Training was conducted with bfloat16 precision to optimize memory usage without compromising accuracy.

Optimization was performed with the 8-bit Paged AdamW optimizer (Loshchilov & Hutter, 2019) implemented in *bitsandbytes* and accessed through Transformers via TrainingArguments(optim="paged_adamw_8bit"). This “paged” variant uses CUDA unified memory to page optimizer states between GPU and host RAM, reducing VRAM requirements while preserving AdamW updates. The final configuration used a learning rate of 2 × 10⁻⁴ and a weight decay of 0.01, selected after a small sweep ({0, 0.01, 0.1}) that showed 0.01 yielded better validation F1 for LoRA-only tuning on our dataset. A cosine learning rate scheduler with a warmup phase constituting 3% of the total training steps was employed to address the sensitivity typically encountered during fine-tuning (Mosbach et al., 2021). A batch size of 4 per device was employed, with gradient accumulation performed over 4 steps, achieving an effective batch size of 16.
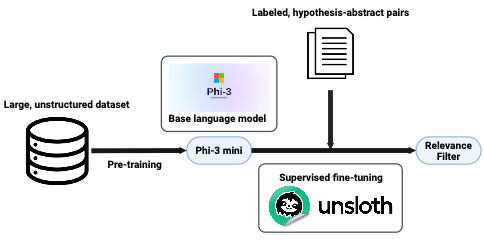

**Figure S2**: Development of the relevance filter using Unsloth for fine-tuning

To train the relevance filter, a dataset comprising 177 abstract-hypothesis pairs was created, consisting of both manually annotated and synthetic examples (see **Generating and Evaluating Synthetic Abstracts** and **Synthetic Abstract Generation - Relevance Filtering Evaluation** sections). Specifically, 140 manually annotated pairs were derived from biomedical literature used in our score alignment evaluation and labeled as "relevant" or "not relevant" for hypothesis evaluation. Additionally, 37 synthetic pairs were created using GPT-4’s API to increase generalization. These abstract-hypothesis pairs adhered to three primary templates: the A-B Hypothesis ("There exists an interaction between the disease {A-term} and the gene {B-term}"), the B-C Hypothesis ("There exists an interaction between the drug {C-term} and the gene {B-term}"), and the A-C Hypothesis ("There exists an interaction between the drug {C-term} and the disease {A-term}"). Each abstract was presented to the model with an instruction to classify it as either 0 (Not Relevant) or 1 (Relevant) in the context of evaluating the provided hypothesis. The dataset was divided into training (105 pairs) and test sets (72 pairs), each with a similar proportion of real and synthetic examples. The misclassifications of the relevance filter on the test set are in the additional file “SKiM_GPT_supplementary_Data.zip”.

**Relevance Filter Prompt Template**

Abstract: {abstract}

Hypothesis: {hypothesis}

Instructions: Classify this abstract as either 0 (Not Relevant) or 1 (Relevant) for evaluating the provided hypothesis.

Score:

###### Synthetic Abstract Generation - Relevance Filtering Evaluation

To generate realistic test cases for evaluating the relevance filter, synthetic biomedical abstracts containing drug–drug interactions (DDIs) and functional gene interactions (FGIs) were created using the GPT-4 API within the previously described Python framework (see **Generating and Evaluating Synthetic Abstracts** section). Each abstract described a specific type of interaction (e.g., "Strong Interaction," "Moderate Interaction," "Direct Physical Interaction," or "Functional Association") or stated the absence of interaction (e.g., "Separate Pathways," "Non-Interacting Proteins," or "Non-Drug Mention") without implying a positive or negative sentiment. For FGIs, experimental methodologies such as yeast two-hybrid screening, co-immunoprecipitation (Co-IP), affinity purification mass spectrometry (AP-MS), surface plasmon resonance (SPR), and nuclear magnetic resonance spectroscopy (NMR) were randomly included to mimic scientific authenticity. Detailed clinical and experimental information, including patient cohorts, study designs, experimental organisms, and statistically supported findings, were incorporated. In this context, categories “Separate Pathways” and “Parallel Functions” are treated as *irrelevant* because the relevance filter’s objective is not to infer mechanistic relationships but to identify whether a text provides usable evidence for evaluating the stated hypothesis (e.g., whether one gene or drug interacts with another). **Supplementary Table T1** provides full definitions and inclusion criteria for all categories.

###### Example Synthetic Abstracts - Relevance Filtering Evaluation

*Relevant to the hypothesis: There exists an interaction between temazepam and trisulfapyrimidines*

PMID: [RANDOM NUMBER] This study aimed to investigate the potential pharmacokinetic and pharmacodynamic interactions between temazepam and trisulfapyrimidines (sulfadiazine).A randomized, double-blind, crossover study was conducted in a cohort of 40 healthy adult volunteers (20 males, 20 females, aged 25-55 years, BMI 18.5-24.9 kg/m2). Participants were randomly assigned to receive either a single oral dose of temazepam (30 mg) and sulfadiazine (500 mg), administered simultaneously, or each drug individually, with a washout period of two weeks between each treatment phase. Blood samples were collected at regular intervals over a 24-hour period following drug administration to determine plasma concentrations of both drugs. Pharmacodynamic parameters, including sleep latency, total sleep time, and number of awakenings, were also assessed using polysomnography.The pharmacokinetic parameters of temazepam and sulfadiazine, including peak plasma concentration (Cmax), time to reach peak plasma concentration (Tmax), and area under the plasma concentration-time curve (AUC), were calculated using non-compartmental analysis. Statistical comparisons were performed using two-way ANOVA, with p<0.05 considered statistically significant. Our results revealed no significant alterations in the pharmacokinetic parameters of temazepam or sulfadiazine when co-administered compared to when administered individually. The Cmax, Tmax, and AUC of both drugs remained unchanged (p>0.05). Similarly, there were no significant differences in the pharmacodynamic parameters of temazepam, including sleep latency, total sleep time, and number of awakenings, between the two treatment phases (p>0.05).In conclusion, this study provides evidence that co-administration of temazepam and sulfadiazine does not result in significant pharmacokinetic or pharmacodynamic interactions. These findings suggest that these drugs can be safely co-administered without the need for dosage adjustments. However, as this study was conducted in healthy volunteers, further research is needed to confirm these findings in patients with underlying health conditions.

*Irrelevant to the hypothesis: There exists an interaction between Potassium Bitartrate and Solriamfetol*

PMID: [RANDOM NUMBER] The present study aims to investigate the physiological effects of Potassium Bitartrate (PB) and Solriamfetol (SL) in a controlled clinical setting.This double-blind, randomized, placebo-controlled trial involved a sample size of 500 adult participants, with an equal distribution of males and females. The subjects were primarily healthy individuals, aged 30-60 years, with no history of chronic diseases. The study was designed to assess the physiological responses to PB and SL, with a focus on cardiovascular and neurological parameters. Participants were randomly assigned to one of three groups: PB (n=166), SL (n=167), and placebo (n=167). The PB group received a daily oral dose of 100mg of PB, the SL group received a daily oral dose of 75mg of SL, and the placebo group received a daily oral dose of a placebo. The intervention period lasted for 12 weeks, with follow-up assessments at 4-week intervals. The primary outcomes measured were heart rate variability (HRV), blood pressure, and sleep latency. Secondary outcomes included cognitive function tests, such as the Montreal Cognitive Assessment (MoCA), and mood assessments using the Hamilton Depression Rating Scale (HDRS). Statistical analysis was performed using one-way ANOVA and post-hoc Tukey's HSD tests. Results indicated a significant decrease in sleep latency in the SL group compared to the placebo group (p<0.01). However, no significant changes were observed in HRV or blood pressure in either the PB or SL group. Cognitive function, as measured by the MoCA, showed no significant differences among the three groups. HDRS scores showed a non-significant trend towards lower scores in the SL group. In conclusion, our findings suggest that Solriamfetol significantly reduces sleep latency, aligning with its known pharmacological profile as a wake-promoting agent. Potassium Bitartrate, on the other hand, did not exhibit significant effects on cardiovascular or neurological parameters in this healthy cohort. These results provide valuable insights into the physiological effects of these substances and further our understanding of their potential clinical applications. Future research should explore these effects in populations with specific health conditions.

**Supplementary Table T1. Category definitions used in the Relevance Filtering Evaluation**

| Dataset | Category | Description | Relevance |
| --- | --- | --- | --- |
| Drug–Drug Interactions (DDI) | Background Information | Mentions of drugs providing only contextual or background detail without direct interaction evidence. | Irrelevant |
| Drug–Drug Interactions (DDI) | Comparative Study | Abstracts comparing drug efficacy or safety without describing co-administration effects. | Irrelevant |
| Drug–Drug Interactions (DDI) | Conflicting Evidence | Reports with contradictory findings regarding interaction presence or severity. | Relevant |
| Drug–Drug Interactions (DDI) | In Vitro Interaction | Experimental results demonstrating or predicting an interaction under controlled laboratory conditions. | Relevant |
| Drug–Drug Interactions (DDI) | Inconclusive | Insufficient or unclear data regarding any pharmacologic or clinical interaction. | Irrelevant |
| Drug–Drug Interactions (DDI) | List Mention | Enumeration of multiple drugs without context linking them mechanistically. | Irrelevant |
| Drug–Drug Interactions (DDI) | Mild Interaction | Describes a modest pharmacokinetic or pharmacodynamic effect with limited clinical significance. | Relevant |
| Drug–Drug Interactions (DDI) | Moderate Interaction | Interaction expected to be clinically relevant and warrant monitoring or dosage adjustment. | Relevant |
| Drug–Drug Interactions (DDI) | Strong Interaction | High-impact or contraindicated co-administration; typically clinically significant. | Relevant |
| Drug–Drug Interactions (DDI) | Potential Interaction | Predictive or hypothesized interaction lacking empirical validation. | Relevant |
| Drug–Drug Interactions (DDI) | Rare Interaction | Uncommon but reported pharmacologic or adverse interaction. | Relevant |
| Drug–Drug Interactions (DDI) | No Significant Interaction | Explicitly reports absence of meaningful pharmacologic effect when co-administered. | Irrelevant |
| Drug–Drug Interactions (DDI) | Non-Drug Mention | Mentions substances not classified as drugs; excluded from DDI scope. | Irrelevant |
| Drug–Drug Interactions (DDI) | Separate Contexts | Drugs discussed in unrelated study contexts with no mechanistic overlap. | Irrelevant |
| Drug–Drug Interactions (DDI) | Variable Interaction | Interaction magnitude or direction varies between studies or populations. | Relevant |
| Functional gene interactions (FGI) | Direct Physical Interaction | Experimental or modeled evidence of direct binding between proteins. | Relevant |
| Functional gene interactions (FGI) | Protein Complex Formation | Co-presence in a multimeric complex without explicit binding characterization. | Relevant |
| Functional gene interactions (FGI) | Enzyme-Substrate Interaction | Catalytic relationship where one protein enzymatically modifies another. | Relevant |
| Functional gene interactions (FGI) | Regulatory Interaction | One protein regulates the expression, localization, or activity of another. | Relevant |
| Functional gene interactions (FGI) | Functional Association | Participation in a shared biological process or function without direct contact. | Relevant |
| Functional gene interactions (FGI) | Genetic Interaction | Phenotypic interaction inferred from genetic perturbation experiments. | Relevant |
| Functional gene interactions (FGI) | Signal Transduction Interaction | Sequential activation or inhibition within signaling cascades. | Relevant |
| Functional gene interactions (FGI) | Protein Modification | Post-translational modification relationship (e.g., phosphorylation, ubiquitination). | Relevant |
| Functional gene interactions (FGI) | Parallel Functions | Proteins perform similar roles in distinct pathways. | Irrelevant |
| Functional gene interactions (FGI) | Separate Pathways | Proteins act in independent biological pathways | Irrelevant |
| Functional gene interactions (FGI) | Different Cellular Compartments | Proteins localized to distinct cellular regions; unlikely to interact directly. | Irrelevant |
| Functional gene interactions (FGI) | No Direct Interaction | Evidence that proteins do not physically associate under tested conditions. | Irrelevant |
| Functional gene interactions (FGI) | Non-Interacting Proteins | Explicitly negative examples from curated databases. | Irrelevant |
| Functional gene interactions (FGI) | Conflicting Evidence | Reports with inconsistent findings on interaction presence or mechanism. | Relevant |

###### Synthetic Abstract Generation - parametric-knowledge leakage experiment

To evaluate SKiM-GPT's susceptibility to information leakage from pre-trained language model parameters, we constructed a set of synthetic biomedical abstracts (see **Generating and Evaluating Synthetic Abstracts)** and tested the system’s ability on abstracts of different sentiments**.** To create abstracts that detail different sentiments (negative, neutral and positive) regarding a given drug and breast cancer, our framework issues prompts to the Deepseek R1 model (temperature: 0.2, max tokens: 2000). Prompts were designed to generate realistic biomedical abstracts following standardized narrative templates and scientific conventions.

To prevent repetition, prompts instructed the model to diversify opening sentences, study types, and outcome measures across abstracts. Randomized selection from a pool of four structural templates (e.g., "Context, Objective, Methods, Results, Conclusion" or "Single-paragraph narrative with embedded methodology") further increased heterogeneity.

The resulting synthetic abstracts were evaluated informally for appropriate technical terminology, logical flow, plausible experimental designs, and conclusions aligned with the desired sentiment.

###### Example Synthetic Abstracts - parametric-knowledge leakage experiment

Positive Sentiment:

PMID: [RANDOM NUMBER] Ado-trastuzumab emtansine (T-DM1) has emerged as a pivotal therapeutic agent for HER2-positive metastatic breast cancer (MBC), demonstrating significant improvements in progression-free survival (PFS) and overall survival (OS) compared to conventional therapies. This study evaluated the efficacy and safety of T-DM1 in a multicenter, randomized phase III trial involving 498 patients with HER2-positive MBC who had previously received trastuzumab and a taxane. Patients were randomized 1:1 to receive either T-DM1 (3.6 mg/kg every 3 weeks) or physician’s choice of lapatinib plus capecitabine (L+C). The primary endpoint was PFS, with secondary endpoints including OS, objective response rate (ORR), and safety. Median PFS was significantly prolonged in the T-DM1 arm (9.6 months vs. 6.4 months; hazard ratio [HR] 0.65, 95% confidence interval [CI] 0.55–0.77, p < 0.001). OS was also markedly improved with T-DM1 (median 30.9 months vs. 25.1 months; HR 0.68, 95% CI 0.55–0.85, p = 0.002). The ORR was higher in the T-DM1 group (43.6% vs. 30.8%, p = 0.008), with a more favorable safety profile, including lower rates of grade ≥3 adverse events (AEs) (40.8% vs. 57.0%, p < 0.001). The most common AEs with T-DM1 were fatigue (36.2%), nausea (33.5%), and thrombocytopenia (28.7%), whereas L+C was associated with higher incidences of diarrhea (58.3%) and hand-foot syndrome (45.6%). These findings underscore T-DM1 as a superior treatment option for HER2-positive MBC, offering clinically meaningful improvements in survival outcomes and tolerability. The data support its integration into standard care regimens for this patient population, particularly after prior trastuzumab failure. Further studies are warranted to explore T-DM1 in earlier lines of therapy and combination strategies.

Neutral Sentiment:

PMID: [RANDOM NUMBER] Alpelisib, a selective phosphatidylinositol 3-kinase (PI3K) inhibitor, has been investigated for its efficacy in treating advanced hormone receptor-positive (HR+), human epidermal growth factor receptor 2-negative (HER2-) breast cancer with PIK3CA mutations. This multicenter, randomized, double-blind, placebo-controlled phase III trial evaluated the addition of alpelisib to fulvestrant in 350 patients with PIK3CA-mutated, HR+/HER2- metastatic breast cancer who had progressed on prior endocrine therapy. The primary endpoint was progression-free survival (PFS), with secondary endpoints including overall survival (OS), objective response rate (ORR), and safety. After a median follow-up of 18.5 months, the median PFS in the alpelisib-fulvestrant arm was 7.4 months (95% CI: 6.9–8.1) compared to 6.9 months (95% CI: 6.2–7.5) in the placebo-fulvestrant arm (hazard ratio [HR] 0.91, 95% CI: 0.75–1.10; p = 0.32). No statistically significant difference in OS was observed (median OS 23.1 vs. 22.8 months; HR 0.97, 95% CI: 0.79–1.19; p = 0.76). The ORR was 21.5% (95% CI: 16.8–26.9) in the alpelisib group versus 18.2% (95% CI: 13.7–23.4) in the placebo group (p = 0.41). Grade 3/4 adverse events, including hyperglycemia (32.1% vs. 2.4%) and rash (15.6% vs. 0.6%), were significantly more frequent with alpelisib. Despite preclinical rationale for PI3K inhibition in PIK3CA-mutated tumors, this study did not demonstrate a clinically meaningful improvement in PFS or OS with alpelisib compared to standard endocrine therapy. These findings suggest that alpelisib may not significantly alter disease progression or survival outcomes in this patient population, highlighting the need for further investigation into predictive biomarkers and combination strategies to optimize PI3K-targeted therapy in breast cancer.

Negative Sentiment:

PMID: [RANDOM NUMBER] Recent clinical trials have investigated the efficacy of abemaciclib, a cyclin-dependent kinase 4/6 (CDK4/6) inhibitor, in the treatment of hormone receptor-positive (HR+), human epidermal growth factor receptor 2-negative (HER2-) advanced breast cancer, with conflicting results regarding its impact on patient outcomes. This multicenter, randomized, double-blind, placebo-controlled phase III study evaluated the effect of abemaciclib in combination with endocrine therapy (letrozole or fulvestrant) versus endocrine therapy alone in 1,248 patients with metastatic HR+/HER2- breast cancer. The primary endpoint was progression-free survival (PFS), with secondary endpoints including overall survival (OS), objective response rate (ORR), and safety. Contrary to prior reports, our findings demonstrated a statistically significant worsening of clinical outcomes in the abemaciclib arm. Median PFS was 9.2 months (95% CI: 8.1–10.3) in the abemaciclib group compared to 12.7 months (95% CI: 11.5–14.0) in the control group (HR: 1.48, 95% CI: 1.25–1.75; p < 0.001). Similarly, median OS was 28.4 months (95% CI: 25.6–31.2) versus 35.1 months (95% CI: 32.3–38.9) in the abemaciclib and control arms, respectively (HR: 1.32, 95% CI: 1.12–1.56; p = 0.003). The ORR was also significantly lower in the abemaciclib group (24.5% vs. 32.8%, p = 0.012). Furthermore, treatment-related adverse events (AEs) were more frequent and severe with abemaciclib, including grade 3/4 neutropenia (38.6% vs. 2.1%), diarrhea (12.4% vs. 1.3%), and fatigue (8.9% vs. 2.7%). These results suggest that abemaciclib, when added to standard endocrine therapy, may detrimentally affect survival and tolerability in this patient population, warranting a reevaluation of its therapeutic role. Further studies are needed to identify subsets of patients who may derive benefit versus harm from this agent.

###### Time-Slicing Parametric-Knowledge Leakage Evaluation

To be able to detect parametric-knowledge leakage (the LLM erroneously relying on its trained weights instead of the retrieved texts to provide a score), the trained weights and the provided texts must disagree in sentiment regarding the hypothesis’ score (*e.g.*, the trained weights alone produce a positive score but the texts indicate a neutral or negative score, or vice versa). This is easiest to accomplish with synthetic abstracts because we can manipulate the text’s sentiment (as discussed in the "Retrieved Text vs Memory” section in the main text of the manuscript). However, it is possible that these synthetic abstracts are different enough from real abstracts that no leakage occurs with synthetic abstracts but does occur with real abstracts. We provide two examples here using real, published abstracts in a time-slicing experiment that indicate the LLM (OpenAI o1 model) bases its evaluation primarily on the texts and not its trained weights.

Prior to 1992, arsenic was well known as a poison and carcinogen. In 1992, arsenic trioxide was discovered as a treatment for acute promyelocytic leukemia (although note that this paper is not in the PubMed corpus) (SUN, 1992). Using PubMed abstracts published in 1992 and earlier, SKiM-GPT scores the hypothesis “Arsenic treats leukemia” with a -2:

*Score: -2 – Reasoning: From the single abstract (PMID: 7023936), arsenic is discussed in the context of its carcinogenic potential and possible role in causing leukemia, rather than treating it. Specifically, it mentions some preliminary indications that arsenic may be linked to lung cancer and leukemia (PMID: 7023936). This contradicts the claim that arsenic treats leukemia, thereby refuting the hypothesis. There is no indication in this abstract of beneficial or therapeutic effects of arsenic on leukemia. Instead, arsenic is implicated as a potential contributing factor to the disease.*

This same hypothesis with no provided texts is scored as a +2:

*Score: +2 – Reasoning: The available abstracts do not provide sufficient details to confirm or deny the hypothesis directly (no PMIDs are cited), so this evaluation relies on general biomedical knowledge. It is well established that arsenic trioxide is used as a standard treatment for a specific subtype of leukemia, namely acute promyelocytic leukemia (APL). In this setting, arsenic promotes the degradation of the disease-driving fusion protein and leads to clinical remissions. Thus, the hypothesis that arsenic can treat leukemia is strongly supported by established therapeutic practice, justifying a score of +2.*

CDK4's role in mammalian cell-cycle progression was characterized in the early-to-mid 1990s. Today, CDK4/6 inhibitors are widely known to treat certain types of breast cancer. Using PubMed abstracts published in 1994 and earlier, SKiM-GPT scores the hypothesis “Inhibiting CDK4 may treat breast cancer” with a +0:

*Score: 0 – Reasoning: The single provided abstract (PMID: 21559680) focuses on how nuclear exclusion of p53 influences CDK2 activity via changes in p21 expression in MCF7 breast cancer cells. While CDK4 is mentioned, the data emphasize that newly synthesized p21 binds CDK2 but not CDK4, and there is no discussion of CDK4 inhibition as a treatment strategy. Consequently, these findings do not provide evidence either for or against the notion that inhibiting CDK4 may treat breast cancer, leaving the hypothesis neither supported nor refuted by the available text.*

This same hypothesis with no provided texts is scored as a +2:

*Score: +2 – Reasoning: The provided abstracts do not contain information on CDK4 and breast cancer, so they are insufficient to evaluate the hypothesis. However, based on general biomedical knowledge, CDK4/6 inhibitors are already in use to treat certain types of breast cancer, particularly hormone receptor-positive subtypes. These therapies have demonstrated clinical benefit and align with the hypothesis that inhibiting CDK4 can help treat breast cancer.*

###### Manual Scoring

To benchmark SKiM-GPT’s score alignment with expert human judgment, we conducted a manual evaluation involving four biomedical researchers who independently scored a set of hypotheses. Each hypothesis represented a unique disease-gene-drug triplet designed to simulate literature-based discovery tasks such as drug repurposing. In total, 14 hypotheses were assessed.

Each evaluator was presented with a structured hypothesis, “{drug} treats {disease} through its effect on {gene}” along with a set of retrieved PubMed abstracts containing co-occurrence of the relevant terms. Two sets of abstracts were evaluated: one unfiltered set derived solely from statistical co-occurrence, and one set of co-occurring abstracts that were filtered using the relevance filter described in the main text.

Evaluators were instructed to assign a score between −2 and +2, where −2 indicated strong refutation of the hypothesis, 0 indicated no support or neutrality, and +2 indicated strong support. Example scoring guidelines are provided below. No consensus discussions were held among evaluators prior to scoring to preserve independence of judgment.

**Scoring Guidelines (AZ - BCHE - 2 PAM)**

Below is an example scoring guideline for the AZ - BCHE - 2PAM SKiM triplet:

*Scoring Guidelines:*

-2: The hypothesis is refuted by consistent evidence indicating that the interactions between Alzheimer's-BCHE and/or BCHE-2 pam contradict the proposed outcome.

-1: The hypothesis is likely refuted based on the evidence. There is moderate indication that the interactions between Alzheimer's-BCHE and/or BCHE-2 pam contradict the proposed outcome, but some uncertainty or contradictory evidence exists.

0: The hypothesis is neither supported nor refuted by the provided texts. The evidence regarding the interactions between Alzheimer's-BCHE and BCHE-2 pam is inconclusive, mixed, lacks sufficient detail, or there is a lack of evidence.

+1: The hypothesis is likely supported by the provided texts. The evidence suggests that the interactions between Alzheimer's-BCHE and BCHE-2 pam may align with the proposed outcome, but some uncertainty or contradictory evidence exists.

+2: The hypothesis is supported by consistent evidence indicating that the interactions between Alzheimer's-BCHE and BCHE-2 pam align with the proposed outcome, with no significant contradictory evidence.

**Example Prompt (AZ - BCHE - 2 PAM)**

*Biomedical Abstracts for Analysis:*

{Retrieved text}

*Assessment Task:*

Evaluate the degree of support for the hypothesis, which posits an interaction between Alzheimer's and 2 pam through their own interactions with BCHE. Use only the provided abstracts from PubMed, each containing only two of the three terms at a time (Alzheimer's + BCHE, or BCHE + 2 pam), to inform your analysis.

*Your evaluation should:*

Integrate evidence from all abstracts to discern how Alzheimer's, BCHE, and 2 pam might be interconnected.

Assess the directionality and nature of each interaction (e.g., activation, inhibition, reactivation) to determine whether it aligns with or contradicts the hypothesis.

Identify and analyze any opposing mechanisms where one interaction may negate the effects of another. Employ logical reasoning, drawing logical inferences where appropriate, based on Alzheimer's-BCHE and BCHE-2 pam interactions.

Assess the nature and directionality of the interactions, determining whether they are beneficial or detrimental to the proposed outcome in the hypothesis.

Be vigilant for any evidence that contradicts or challenges the hypothesis, explicitly addressing any contradictions in your reasoning.

Avoid inferring effects not supported by the texts, ensuring that all conclusions are grounded in the provided information.

Ensure that your scoring reflects an unbiased assessment based solely on the provided evidence, considering logical inferences from indirect evidence.

*Examples:*

Example 1: Strong Positive Outcome - Hypothesis: 2 pam treats Alzheimer's through BCHE. - Evidence: - Multiple abstracts show that 2 pam inhibits BCHE. - Inhibition of BCHE significantly improves Alzheimer's in various contexts. - Logical Conclusion: - Since 2 pam inhibits BCHE, and inhibition of BCHE improves Alzheimer's, there is strong, consistent indirect evidence supporting the hypothesis. - Scoring: - The interactions are consistent and beneficial. Assigned Score: +2

Example 2: Likely Positive Outcome - Hypothesis: 2 pam treats Alzheimer's through BCHE. - Evidence: - Some abstracts suggest that 2 pam may activate BCHE. - Activation of BCHE might improve Alzheimer's, but evidence is limited or not robust. - Logical Conclusion: - There is evidence indicating potential beneficial interactions, but some uncertainty exists due to limited data or minor contradictions. - Scoring: - The hypothesis is likely supported by the evidence. - Assigned Score: +1

Example 3: Neutral Outcome - Hypothesis: 2 pam treats Alzheimer's through BCHE. - Evidence: - Abstracts provide insufficient or inconclusive information about the interactions. Evidence might be mixed or does not directly relate to the hypothesis. - - Logical Conclusion: There is not enough evidence to support or refute the hypothesis. - Scoring: - The evidence is inconclusive. - Assigned Score: 0

Example 4: Likely Negative Outcome - Hypothesis: 2 pam treats Alzheimer's through BCHE. - Evidence: - Some abstracts indicate that 2 pam activates BCHE. - Activation of BCHE may worsen Alzheimer's, but evidence is limited or not definitive. - Logical Conclusion: - There is moderate indication that the interactions may be detrimental to the proposed outcome, but some uncertainty or exceptions exist. - Scoring: - The hypothesis is likely refuted based on the evidence. - Assigned Score: -1-

Example 5: Strong Negative Outcome - Hypothesis: 2 pam treats Alzheimer's through BCHE. - Evidence: - Multiple abstracts consistently show that 2 pam inhibits BCHE. - Inhibition of BCHE clearly worsens Alzheimer's across various studies. - No evidence suggests any interaction aligns with supporting the hypothesis. - Logical Conclusion: - Since 2 pam inhibits BCHE, and inhibition of BCHE worsens Alzheimer's, the hypothesis is clearly refuted by strong, consistent evidence. - Scoring: - The interactions do not align with supporting the hypothesis. Assigned Score: -2

Your goal is to determine the degree of support for the hypothesis:

*2 pam treats Alzheimer's through its effect on BCHE.*

*Instructions:*

1. Review each abstract to understand how Alzheimer's, BCHE, and 2 pam might be interconnected based on the available information.

2. Identify and assess the nature of each interaction between the terms (e.g., activation, inhibition, reactivation).

3. Determine whether each interaction aligns with or contradicts the hypothesis based on its nature:

Alignment: Interactions that support the hypothesis (e.g., inhibition of BCHE by 2 pam leading to improvement in Alzheimer's).

Contradiction: Interactions that oppose the hypothesis (e.g., activation or reactivation of BCHE by 2 pam leading to worsening of Alzheimer's).

4. Synthesize the findings from multiple abstracts, considering how the interactions fit together to support or refute the hypothesis: 2 pam treats Alzheimer's through its effect on BCHE.

5. Provide a justification for your scoring decision based on the analysis. Explain your reasoning step-by-step in terms understandable to an undergraduate biochemist.

Focus on:

Explaining the logical connections and the directionality of relationships.

Determining whether the interactions support or contradict the hypothesis, using indirect evidence and logical inferences.

Assessing if the interactions are beneficial or detrimental to the proposed outcome.

6. Be vigilant for any evidence that contradicts or challenges the hypothesis. Address any contradictions explicitly in your reasoning.

7. Verify any assumptions about the roles and effects of Alzheimer's, BCHE, and 2 pam as presented in the abstracts. Avoid inferring effects not supported by the texts.8. In your final assessment, explicitly cite the scoring guideline that corresponds to your conclusion. Explain why the evidence meets the criteria for that specific score.

Note: Pay close attention to the nature of the interactions between entities. An interaction that activates a harmful process may be detrimental, while inhibition of a beneficial process may also be detrimental. Consider whether such interactions contradict the hypothesis.

*Definitions:*

Evidence: Multiple sources agree and provide clear indications supporting a particular interaction and its effects.

Contradictory Evidence: Evidence that directly opposes the hypothesis, showing that the proposed mechanism of action does not produce the expected outcome.

Directionality of Interaction: The specific effect an interaction has (e.g., activation, inhibition, reactivation) and whether it supports or opposes the hypothesis.

*Checklist Before Finalizing Your Response:*

Have you addressed whether the interactions support or contradict the hypothesis?- Have you assessed if the interactions are beneficial or detrimental?

Have you explicitly cited the scoring guideline that matches your conclusion?

Have you explained why the evidence meets the criteria for the assigned score?

Format your response as: Score: [Number] Point(s) - Reasoning: [Reasoning]

**LBD False-Positive Removal Evaluation Details**

SKiM was run with the following parameters. The A term was “breast cancer”. The B terms were a list of human genes. The C terms were a list of FDA-approved drugs. The two term lists are in the additional file “SKiM_GPT_supplementary_Data.zip”.

###### How to Identify Novel Hypotheses Returned by SKiM-GPT

The SKiM-GPT system evaluates many A-B-C hypotheses. Many of these links and their associated hypotheses are likely to be already established in the literature. Known links/hypotheses can be identified because the direct A-C link (for instance between the drug and disease in our drug-gene-disease example) is also strong by co-occurrence and is strongly supported by LLM analysis, and therefore already likely known. The known A-C and A-B-C relationships are not necessarily problematic, as it validates the ability of SKiM-GPT to find known relations, and they are easily filtered by the presence of the A-C direct link. In contrast, a strong A-B-C link with a weak or absent A-C link can be used to find relationships that are likely novel. In our list of 14 disease-gene-drug relationships (**Figure 3**), we have 2 examples of links that are likely novel. These are the A-B-C links of: 1) Alzheimer’s - FYN - pp2 and 2) Non-alcoholic fatty liver disease - ULK1 - pp242 (where FYN and ULK1 are genes linking disease to drug). The corresponding A-C (disease-drug) direct links both have model scores of 0, suggesting a lack of literature support for these drugs as known treatments for the associated disease. However, the A-B-C links have a positive model score (+1) indicating that the drugs may affect the disease in a beneficial way through the interaction with the associated gene. Thus, these A-B-C links form novel hypotheses that could be tested pre-clinically.

**Supplementary Table T2:** Table of scores for A-C and A-B-C disease/drug relationships

| A-C Relationship | SKiM-GPT AC Score Median | A-B-C Relationship | SKiM-GPT ABC Score Median |
| --- | --- | --- | --- |
| Alzheimer's - 2 pam | -2 | Alzheimer's - BCHE - 2 pam | -2 |
| Alzheimer's - pp2 | 0 | Alzheimer's - FYN - pp2 | 1 |
| Breast cancer - abemaciclib | 2 | Breast cancer - CDK4 - abemaciclib | 2 |
| Breast cancer - estrogens | -2 | Breast cancer - ESR1 - estrogens | -2 |
| Diabetes - stavudine | -2 | Diabetes - AHR - stavudine | 0 |
| Diabetes - tirzepatide | 2 | Diabetes - GIPR - tirzepatide | 2 |
| Diabetes - zileuton | -1 | Diabetes - LOX - zileuton | 1 |
| Heart disease - erlotinib | 1 | Heart disease - EGFR - erlotinib | 2 |
| Heart disease - gefitinib | 1 | Heart disease - RAF1 - gefitinib | 1 |
| Lupus - isoniazid | -2 | Lupus - NAT2 - isoniazid | -1 |
| Non-alcoholic fatty liver disease - 6-formylindolo(3,2-b)carbazole | 0 | Non-alcoholic fatty liver disease - AHR - 6-formylindolo(3,2-b)carbazole | -0.5 |
| Non-alcoholic fatty liver disease - pp242 | 0 | Non-alcoholic fatty liver disease - ULK1 - pp242 | 1 |
| Pancreatic cancer - gant61 | 2 | Pancreatic cancer - CCK - gant61 | 0 |
| Pancreatic cancer - ginsenoside rb1 | 1 | Pancreatic cancer - NAMPT - ginsenoside rb1 | -2 |

###### Exact String Matches and Evaluating Mechanisms

The SKiM search algorithm searches by exact text matches but allows “and” (&) and “or” (|) logical operators. This allows searching for mechanisms such as listing multiple pathways or biological processes as the B term to explore how two genes might interact or how a drug might treat a disease. For example, in the original SKiM paper, we were able to detect the same interactions discovered by Swanson in 1986 where fish oil was found to be an effective treatment for Raynaud’s disease based on its effect on blood viscosity (Millikin et al., 2023; Swanson, Don R., 1987). If the user wanted to explore potential mechanisms linking fish oil to Raynaud’s disease, they could run SKiM with an A term of “Raynaud’s”, a C term of “Fish Oil”, and then use a list of more than 23,000 bio-processes (including blood viscosity and other terms) as the B terms. One hypothesis template could be “Fish oil treats Raynaud’s via its action on {B-Term}”. When SKiM is run using these terms and hypotheses and censored to 1985 and before (prior to Swanson’s discovery), three B terms are significant (with a *p*-value < 1e-5): “platelet aggregation”, “aggregation”, and “blood viscosity”, all of which are related to blood viscosity. When the relevant abstract sets are analyzed by SKiM-GPT all three return a score of +1, indicating that the hypotheses are likely supported (**Figure S3**).

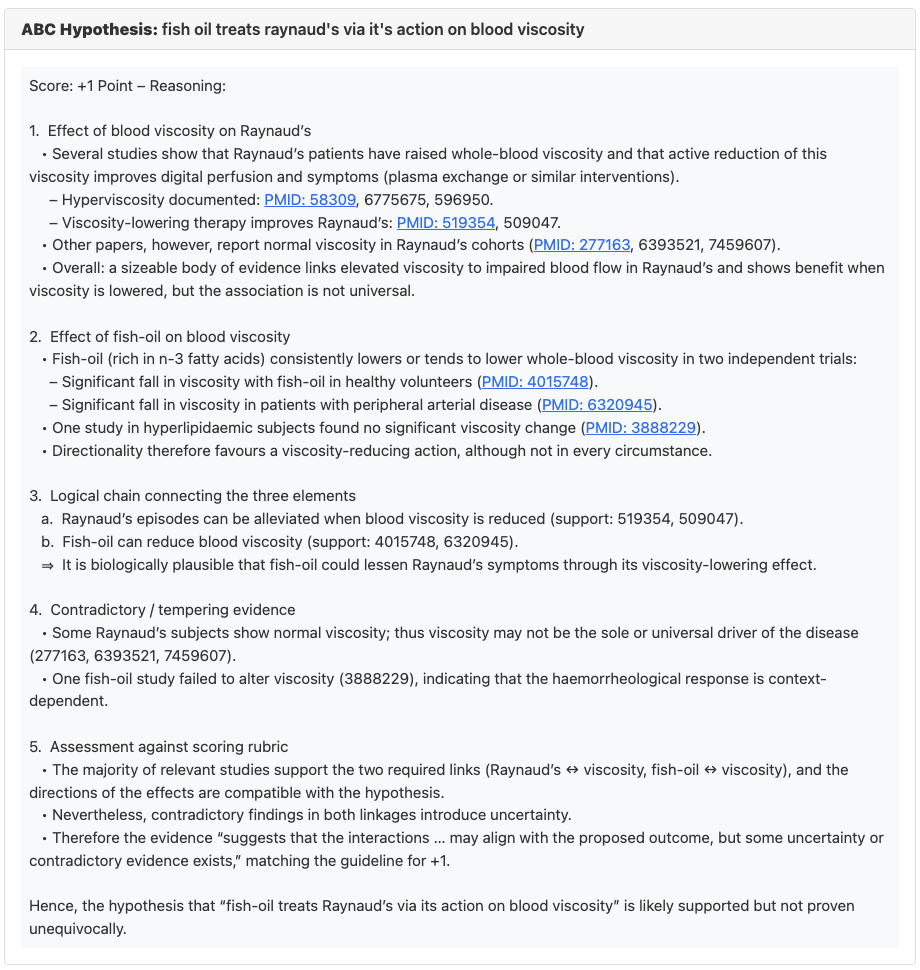

**Figure S3:** Score and rationale for hypothesis “Fish oil treats Raynaud’s via its action on blood viscosity”**.**
